# DeepGOMeta: Predicting functions for microbes

**DOI:** 10.1101/2024.01.28.577602

**Authors:** Rund Tawfiq, Kexin Niu, Robert Hoehndorf, Maxat Kulmanov

## Abstract

Analyzing microbial samples remains computationally challenging due to their diversity and complexity. The lack of robust *de novo* protein function prediction methods exacerbates the difficulty in deriving functional insights from these samples. Traditional prediction methods, dependent on homology and sequence similarity, often fail to predict functions for novel proteins and proteins without known homologs. Moreover, most of these methods have been trained on largely eukaryotic data, and have not been evaluated or applied to microbial datasets. This research introduces DeepGOMeta, a deep learning model designed for protein function prediction, as Gene Ontology (GO) terms, trained on a dataset relevant to microbes. The model is validated using novel evaluation strategies and applied to diverse microbial datasets. Data and code are available at https://github.com/bio-ontology-research-group/deepgometa

## Introduction

Protein function prediction has evolved significantly over the past few years, transitioning from reliance on basic sequence alignment to approaches based on machine learning, natural language processing, or analysis of biological networks [1]. Despite these advances, few methods have been developed for and evaluated on metagenome or amplicon sequencing data mainly because there is no “ground truth” unless the methods are applied to “mock communities” which are highly simplified versions of actual microbial communities and not representative of the complexities encountered in real-world cases.

Microbial communities are especially complex, mainly due to the diversity of organisms they contain, including many that have yet to be cultured. This gives rise to a phenomenon often termed metagenomic ‘dark matter’ where 50% to 80% of metagenomic proteins remain unannotated using current methods [2]. This complexity and diversity often renders traditional annotation methods inadequate, particularly when presented with novel proteins.

Microbial genomic data primarily come in two forms: amplicon sequences and whole genome sequencing (WGS) reads. Amplicon sequences, like 16S rRNA, are key for bacterial taxonomic classification, but their utility in function prediction is limited. Tools like PICRUSt2 [3] and Tax4Fun2 [4] infer microbial community functions using homology-based algorithms by aligning to reference databases. However, the accuracy of these predictions is constrained by algorithm limitations and database completeness. WGS enables the reconstruction of complete microbial genomes, allowing for a more direct assessment of a microbial community’s functional potential, traditionally done by aligning protein-coding sequences to known proteins using algorithms like BLAST [5]. Existing protein function prediction methods face significant limitations in microbial contexts. Even when enhanced with machine learning, these methods are limited by their training datasets. For example, the Critical Assessment of Function Annotation (CAFA) challenge [6] utilizes the SwissProt database, rich in eukaryotic proteins, overlooking the predominantly prokaryotic nature of metagenomes [7]. Moreover, most of these methods have not been validated on or applied to microbial data, largely due to the lack of robust evaluation strategies. These limitations highlight the need for models trained on relevant data and innovative evaluation strategies.

Deep learning has shown remarkable potential in analyzing biological data through its ability to detect intricate patterns in vast datasets [8]. DeepGOMeta incorporates ESM2 (Evolutionary Scale Modeling 2) [9], a deep learning framework that extracts meaningful features from protein sequences by learning from evolutionary data. By utilizing these learned features through ESM2, and training on a more representative dataset, DeepGOMeta can predict protein functions even in the absence of explicit sequence similarity or homology to known proteins. Moreover, we introduce novel evaluation strategies to assess the method’s performance when applied to microbial data. Taken together, DeepGOMeta addresses the multifaceted challenges associated with protein function prediction for microbial data.

## Methods

### Materials and Data

#### UniProtKB/Swiss-Prot Dataset and Gene Ontology

We obtained all proteins that were manually curated and reviewed from the UniProtKB/Swiss-Prot Knowledgebase (v2023 03, r28-June-2023) [10]. We further filtered to select for proteins that belong to prokaryotes, archaea and phages, and only kept proteins with experimental functional annotations using evidence codes EXP, IDA, IPI, IMP, IGI, IEP, TAS, IC, HTP, HDA, HMP, HGI, HEP. The dataset contains 10, 107 reviewed and manually annotated proteins.

Metagenomes contain many uncharacterized, novel proteins and in order to evaluate our models on novel proteins, we generated training, validation and testing splits based on sequence similarity. First, we grouped the proteins by their similarity using Diamond (v2.0.9) [11] (e-value 0.001) and split them into training, validation and testing sets, 81/9/10 %, respectively. This is to ensure that the training and validation set proteins do not have any similar sequences in the training set.

We trained and evaluated a model for each of the GO sub-ontologies separately (r2023-01-01) [12]. Table 1 summarizes the datasets for each sub-ontology.

**Table 1.** Summary of the UniProtKB/Swiss-Prot dataset.

| Ontology | Terms | Proteins | Groups | Train | Valid | Test | Time |
| --- | --- | --- | --- | --- | --- | --- | --- |
| MFO | 3,022 | 6,280 | 2,264 | 5,297 | 354 | 629 | 30 |
| BPO | 4,861 | 6,620 | 2,717 | 5,563 | 453 | 604 | 45 |
| CCO | 537 | 4,975 | 2,539 | 4,174 | 398 | 403 | 26 |

The table shows the number of GO terms, number of proteins in training, validation, and testing sets for the UniProtKB/Swiss-Prot dataset.

To compare our model against other methods, we generated a test set by following the CAFA [6] challenge time-based approach. We downloaded UniProtKB/Swiss-Prot (v2023 05 r2023-11-08) and extracted newly annotated proteins in this version.

### Protein-Protein Interactions data

For the 10,107 proteins in our dataset, we obtained protein– protein interaction (PPI) data from the STRING (v11.0) [13] database, which yielded 14,524 interactions. There were 7 different modes of interactions: binding, activation, reaction, catalysis, expression, inhibition, and ptmod.

### Microbial Benchmark Datasets

#### MGnify dataset

To evaluate microbial protein annotations, we downloaded the MGnify protein database (r2023 02) [14] and its associated metadata. This database includes protein sequences from publicly available metagenomic assemblies within MGnify. We extracted 2,000 random proteins from this database, where half were ‘aquatic’ and half were ‘terrestrial’ (lineage:root:Environmental:Terrestrial/Aquatic).

#### Paired 16S and WGS dataset

We applied our method to generate functional profiles of microbial communities using four publicly available datasets that contain both 16S amplicon data and WGS from the same samples, shown in Table 2. We used two human stool microbiomes: 60 samples from Indian individuals (PRJNA397112) and 60 samples from Cameroonian individuals (WGS: PRJEB27005, 16S: mgp15238) [15, 16], an environmental microbiome: 22 blueberry plant soil samples (WGS: PRJNA484230, 16S: PRJNA389786), and 11 mammalian stool samples (WGS: SRP115632, 16S: SRP115643). The datasets represent a variety of host-associated and environmental microbiomes.

**Table 2.** Descriptions of the paired datasets used for evaluation.

| Name | Biome | N | Region | OTUs |
| --- | --- | --- | --- | --- |
| India | human stool | 60 | V3 | 2077 |
| Cameroon | human stool | 60 | V5-V6 | 721 |
| Blueberry | terrestrial | 22 | V6-V8 | 1824 |
| Mammalian Stool | mammalian stool | 11 | V6-V8 | 420 |

Biome, number of samples, 16S rRNA gene region and number of identified OTUs for each dataset.

### Baseline and Comparison methods

For our evaluations, we used baseline methods that do not rely on predictions based on sequence similarity, as our aim is to test the predictors on challenging sequences. Therefore, we do not include methods that are primarily based on sequence similarity, such as BLAST, Diamond, or their combinations, as baselines. For the time-based dataset evaluation, we selected three state-of-the-art methods developed by other groups: [17], SPROF [18] and NetGO3 [19].

#### Naive approach

Due to the imbalance in GO class annotations and propagation based on the true-path-rule, some classes have more annotations than others. Therefore, it is possible to obtain prediction results just by assigning the same GO classes to all proteins based on annotation frequencies. In order to test the performance obtained based on annotation frequencies, CAFA introduced a baseline approach called “naive” classifier [6]. Here, each query protein *p* is annotated with GO classes with a prediction scores computed as:

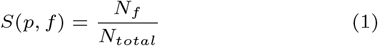

where *f* is a GO class, *N*_*f*_ is a number of training proteins annotated by GO class *f*, and *N*_*total*_ is a total number of training proteins. We implemented the same method.

#### MLP (ESM2)

The MLP baseline method predicts protein functions using a multi-layer perception (MLP) from a protein’s ESM2 embedding [9]. We generated an embedding vector of size 5,192 using the ESM2 15B model and passed it to two layers of MLP blocks where the output of the second MLP block had residual connection to the first b lock. T his r epresentation i s p assed to the final c lassification la yer wi th si gmoid ac tivation function. One MLP block performs the following operations:

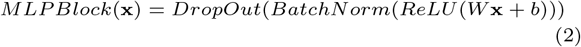

The input vector **x** of length 5, 192 represents the ESM2 embedding and is reduced to 1, 024 by the first MLPBlock:

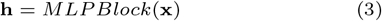

This representation is passed to the second MLPBlock with the input and output size of 1, 024 and added to itself using a residual connection:

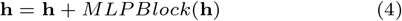

Finally, we passed this vector to a classification layer with a sigmoid activation function. The output size of this layer is equal to the number of classes in each sub-ontology:

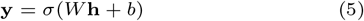

We trained a different model for each sub-ontology in GO.

#### DeepGO-PLUS and DeepGOCNN

DeepGO-PLUS [20] predicts protein functions by combining DeepGOCNN, which predicts functions from the amino acid sequence of a protein using a 1-dimensional convolutional neural network (CNN), and the DiamondScore method. DeepGOCNN captures sequence motifs that are related to GO functions. Here, we only used CNN based predictions.

#### DeepGOZero

DeepGOZero [21] combines protein function prediction with a model-theoretic approach for embedding ontologies into a distributed space, ELEmbeddings [22]. ELEmbeddings represent classes as *n*-balls and relations as vectors to embed ontology semantics into a geometric model. It uses InterPro domain annotations represented as binary vector as input and applies two layers of MLPBlock as in our MLP baseline method to generate an embedding of size 1024 for a protein. It learns the embedding space for GO classes using ELEmbeddings loss functions and optimizes together with protein function prediction loss. For a given protein *p* DeepGOZero predicts annotations for a class *c* using the following formula:

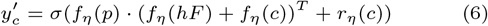

where *f*_*η*_ is an embedding function, *hF* is the hasFunction relation, *r*_*η*_(*c*) is the radius of an *n*-ball for a class *c* and *σ* is a sigmoid activation function. It optimizes binary crossentropy loss between predictions and the labels together with ontology axioms losses from ELEmbeddings.

#### TALE

TALE [17] predicts functions using a transformer-based deep neural network model which incorporates hierarchical relations from the GO into the model’s loss function. The deep neural network predictions are combined with predictions based on sequence similarity. We used the trained models provided by the authors to evaluate them in the time-based dataset.

#### SPROF-GO

SPROF-GO [18] method uses the ProtT5-XL-U50 [23] protein language model to extract proteins sequence embeddings and learns an attention-based neural network model. The model incorporates the hierarchical structure of GO into the neural network and predicts functions that are consistent with hierarchical relations of GO classes. Furthermore, SPROF-GO combines sequence similarity-based predictions using a homology-based label diffusion algorithm. We used the trained models provided by the authors to evaluate them on the time-based dataset.

#### Pathway prediction

PICRUSt2 [3] provides the potential functions of microbial communities using 16s rRNA data and a reference genome databases. We used operational taxonomic unit (OTU) tables as the input for PICRUSt2 and focused on MetaCyc [24] pathways and their abundance scores. We performed Principal Component Analysis (PCA) and k-means clustering to discern patterns within the dataset based on these MetaCyc pathway features. The value of k was determined based on the number of categories within each phenotype. We measured clustering purity based on the true phenotype labels in the datasets (eq. 16).

### Evaluation

#### Evaluation metrics

We used four different measures to evaluate the performance of our models. Three protein-centric measures *F*_max_, *S*_min_ and AUPR and one class-centric AUC.

*F*_max_ is a maximum protein-centric F-measure computed over all prediction thresholds. First, we computed average precision and recall using the following formulas:

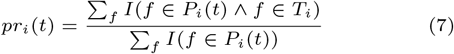

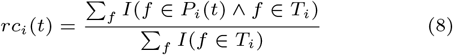

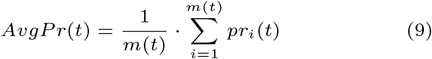

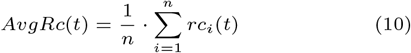

where *f* is a GO class, *T*_*i*_ is a set of true annotations, *P*_*i*_(*t*) is a set of predicted annotations for a protein *i* and threshold *t, m*(*t*) is a number of proteins for which we predict at least one class, *n* is a total number of proteins and *I* is an indicator function which returns 1 if the condition is true and 0 otherwise. Then, we compute the *F*_max_ for prediction thresholds *t* ∈ [0, 1] with a step size of 0.01. We count a class as a prediction if its prediction score is greater than or equal to *t*:

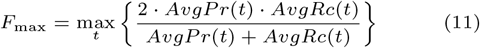

*S*_min_ computes the semantic distance between real and predicted annotations based on information content of the classes. The information content *IC*(*c*) is computed based on the annotation probability of the class *c*:

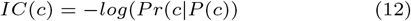

where *P* (*c*) is a set of parent classes of the class *c*. The *S*_min_ is computed using the following formulas:

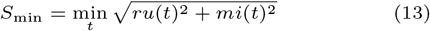

where *ru*(*t*) is the average remaining uncertainty and *mi*(*t*) is average misinformation:

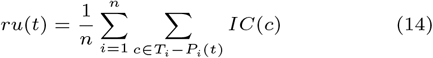

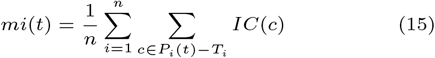

AUPR is the area under the average precision (*AvgPr*) and recall (*AvgRc*) curve.

AUC is a class-centric measure where we computed AUC ROC per class and calculated the average.

Purity assesses the homogeneity of clusters formed by a *k*-means clustering algorithm. We clustered samples based on their predicted functions and used purity to evaluate whether the samples with same phenotype are in the same cluster. The Weighted Average Clustering Purity (WACP) formula is given by:

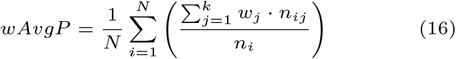

where *N* is the total number of data points, *k* is the number of clusters, *n*_*ij*_ is the number of data points from cluster *j* that are assigned to cluster *i, n*_*i*_ is the total number of data points assigned to cluster *i*, and *w*_*j*_ is the weight associated with cluster *j*.

We calculated function abundance to provide a quantitative assessment of the functional potential within a microbial sample. The abundance of a function (*A*(*f*) is the sum of the relative abundance of all taxa present in a sample that contain a certain function, given by:

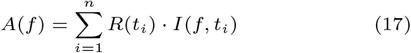

where *i* is an index representing each taxon, *n* is the total number of taxa in the sample, *R*(*t*_*i*_) is the relative abundance of the *i*^*th*^ taxon, and *I*(*f, t*_*i*_) is an indicator function that equals 1 if the *i*^*th*^ taxon contains function *f*, and 0 otherwise.

#### MGnify dataset

We annotated the 2,000 randomly selected proteins with DeepGOMeta, which annotates each protein with GO terms. We used two clustering approaches for the first evaluation. The first approach, sequence similarity clustering, involved calculating pairwise sequence similarities between the proteins using DIAMOND BLASTp (v2.1.8) [11], followed by dimensional reduction using t-Distributed Stochastic Neighbor Embedding (t-SNE), and k-means clustering with k=2 based on the binary nature of the dataset’s phenotypes. We calculated clustering purity using the known environment labels of the proteins.

For the second approach, semantic similarity clustering, we filtered the GO annotations resulting from DeepGOMeta to retain the most specific terms for each protein. For measuring the semantic similarity between protein pairs, we utilized Resnik’s similarity method [25], combined with Best Match Average (BMA) strategy. Resnik’s similarity measure is defined as the most informative common ancestor (MICA) of the compared classes in the ontology. First, we computed information content (IC) for every class with following formula:

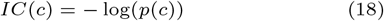

Then, we found Resnik’s similarity by:

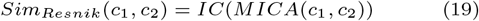

We computed all possible pairwise similarities of two annotation sets and combined them with:

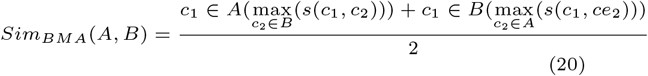

where *s*(*x, y*) = *Sim*_*Resnik*_(*x, y*).

We then performed a similar dimensionality reduction and clustering using t-SNE, k-means and a purity calculation (eq. 16).

We further subsetted the 2,000-protein Mgnify dataset to only keep proteins with existing Pfam annotations (n = 567) [26]. For these proteins, we used Pfam2GO to map Pfam and GO annotations. We calculated purity using the same semantic similarity clustering approach described earlier.

#### Paired dataset

We analyzed four diverse microbiome datasets, each containing paired 16S rRNA amplicon and WGS data. For the 16S data, we used a Nextflow pipeline employing the RDP classifier (v18) for processing and taxonomic classification available on our GitHub repository^1^. We sourced protein sequences corresponding to the identified bacteria in the RDP database from NCBI and annotated with DeepGOMeta [27, 28]. We then constructed functional profiles for each sample by aggregating the DeepGOMeta-derived functions of all bacteria present, weighing each function by the relative abundance of the genera in which it was present (eq. 17). We also constructed a a binary matrix of all the samples and functions in the dataset, where the presence of a function in a sample is represented by 1 and the absence by 0.

For WGS data, we used fastp (v0.23.2) [29] for trimming [-q 30]. For host-associated microbiome samples, we used Bowtie2 (v2.5.1) [30] to filter out reads mapping to the host’s reference genome. We then assembled the reads with MEGAHIT (v1.2.9) [31], predicted protein sequences with prodigal (v2.6.3) [32], and annotated the predicted proteins using DeepGOMeta. For each sample, we constructed a functional profile by aggregating the functions derived from DeepGOMeta annotations of all proteins present in the sample. We constructed a binary matrix for these results as described above.

For each dataset, we applied PCA and *k*-means clustering to the OTU table containing the relative abundance of bacterial genera. The choice of *k* in *k*-means clustering was determined by the number of phenotype categories present for each phenotype under investigation. We calculated clustering purity based on the known phenotype category labels provided in the metadata (eq. 16). We conducted this analysis for all categorical phenotypes across each dataset.

## Results

### DeepGOMeta

Microbial samples are complex and contain many uncharacterized proteins. Previously, we developed DeepGO-SE [33], a method for protein function prediction using protein sequence embeddings generated by ESM2 [9] and approximate semantic entailment. We showed that DeepGO-SE can be applied to uncharacterized proteins; however, since it is trained on all experimentally annotated proteins form UniProt-KB/Swissprot database, many of the functions it predicts are not relevant to microbiomes and exist only in eukaryotic genomes. Therefore, we trained DeepGOMeta, a specific version of DeepGO-SE, optimized to predict functions of organisms found in microbiomes. We created a dataset of prokaryotic, archaeal and viral proteins with experimental annotations from UniProt-KB/SwissProt and trained and evaluated three models for the three sub-ontologies of GO. In addition, we created a time-based benchmark dataset in order to compare with DeepGO-SE and other state-of-the-art function prediction methods.

Proteins do not function in isolation and PPIs play significant role in biological processes that take place in the environment. PPI networks also offer a means to reveal functional information for unknown proteins within microbial datasets. In order to test if PPIs help to improve protein function predictions, we trained a model which combines PPIs from STRING Database [13] using Graph Attention Networks. We refer to this model as DeepGOMeta-PPI.

We developed novel evaluation strategies to test the performance of DeepGOMeta in annotating proteins derived from microbial data, and we used these strategies to test the method against sequence-similarity clustering and Pfam database annotations. We also developed two different workflows for functional characterization of microbial samples consisting of 16S amplicon and WGS reads. In the case of 16S amplicon reads, we use OTUs to predict functions by utilizing the reference genomes of the genera in the samples.We then aggregate all the functions that were annotated into a functional profile for that genus. In the case of WGS reads, we performed *de novo* metagenome assembly and predicted functions from metagenome assemblies. Figure 1 depicts these workflows. We applied DeepGOMeta to diverse microbial datasets, and compared functional profiles, pathways, and taxonomy-based methods to gain biological insights.

**Fig. 1.**
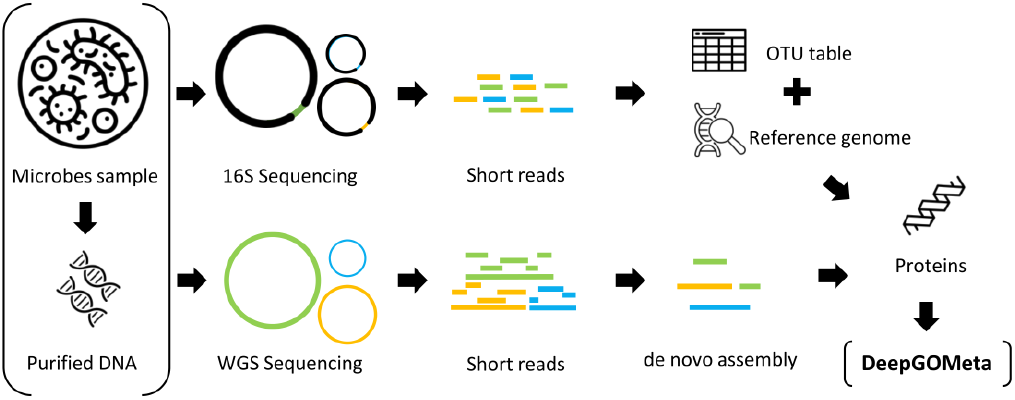
The figure provides an overview of the workflows used to generate functional profiles using DeepGOMeta for amplicon samples and WGS samples.

### Evaluation on the similarity-based benchmark

We trained, validated and tested our models for the three sub-ontologies of GO using the UniProtKB/Swiss-Prot dataset splitted based on sequence similarity (See Methods section). We compared with four baseline methods such as MLP(ESM2), DeepGraphGO, InterPro and Naive. We selected these methods because they do not rely on sequence similarity to predict functions.

In the MFO evaluation, DeepGOMeta performed best in all evaluation metrics. It performed slightly better than MLP(ESM2) in terms of *F*_max_ and *S*_min_,; however, the AUPR and term-centric AUC were significantly better. Combining PPI network features into the model reduced its performance, but was still better than the DeepGraphGO method, which is also based on PPIs. Table 3 provides the evaluation results for MFO classes.

**Table 3.** Evaluation results for Molecular Function Ontology classes.

| Method | $F_{\max}$ | $S_{\min}$ | AUPR | AUC |
| --- | --- | --- | --- | --- |
| MLP(ESM2) | 0.465 | 12.587 | 0.432 | 0.723 |
| DeepGOMeta | **0.468** | **12.230** | **0.449** | **0.874** |
| DeepGOMeta-PPI | 0.436 | 12.909 | 0.392 | 0.839 |
| DeepGraphGO | 0.315 | 14.385 | 0.223 | 0.456 |
| InterPro | 0.208 | 13.685 | 0.308 | 0.588 |
| Naive | 0.312 | 14.398 | 0.161 | 0.500 |

This table shows protein-centric $F_{\max}$ , $S_{\min}$ , and AUPR, and the class-centric average AUC.

In the BPO evaluation, our model resulted in best *F*_max_ of 0.476 which was significantly better (Wilcoxon signed-rank test p-value is 8 *·* 10^−37^) than the second best MLP(ESM2) baseline. Combining PPI networks in DeepGOMeta did not improve the model and lead to slightly lower *F*_max_ of 0.469. Interestingly, InterPro baseline performance was close to *F*_max_ of 0. We believe that this might be due to the fact that not many of InterPro annotations are linked to BPO classes. It also explains the low performance of the DeepGraphGO method which uses InterPro annotations. Table 4 provides detailed evaluation results.

**Table 4.** Evaluation results for Biological Process Ontology classes.

| Method | $F_{\max}$ | $S_{\min}$ | AUPR | AUC |
| --- | --- | --- | --- | --- |
| MLP(ESM2) | 0.449 | 20.095 | 0.419 | 0.811 |
| DeepGOMeta | **0.476** | **19.394** | **0.462** | **0.847** |
| DeepGOMeta-PPI | 0.469 | 19.715 | 0.443 | 0.823 |
| DeepGraphGO | 0.340 | 22.607 | 0.283 | 0.507 |
| InterPro | 0.040 | 24.691 | 0.294 | 0.519 |
| Naive | 0.337 | 22.564 | 0.248 | 0.500 |

This table shows protein-centric $F_{\max}$ , $S_{\min}$ , and AUPR, and the class-centric average AUC.

In the CCO evaluation, DeepGOMeta achieved the best *F*_max_ of 0.739 followed by almost the same performance by MLP(ESM2) baseline. Noticably, MLP(ESM2) method resulted in the best *S*_min_. Similarly to MFO and BPO evaluations, combining PPIs did not improve the predictions. DeepGraphGO method resulted in *F*_max_ of 0.501 which is slightly better than Naive classifier, and InterPro annotation-based prediction performance was close to zero. Table 5 provides the evaluation results.

**Table 5.** Evaluation results for Cellular Component Ontology classes.

| Method | $F_{\max}$ | $S_{\min}$ | AUPR | AUC |
| --- | --- | --- | --- | --- |
| MLP(ESM2) | 0.738 | **3.369** | 0.741 | 0.897 |
| DeepGOMeta | **0.739** | 3.394 | **0.759** | **0.903** |
| DeepGOMeta-PPI | 0.706 | 3.648 | 0.710 | 0.869 |
| DeepGraphGO | 0.501 | 5.218 | 0.494 | 0.577 |
| Interpro | 0.075 | 6.067 | 0.322 | 0.539 |
| Naive | 0.498 | 5.367 | 0.308 | 0.500 |

This table shows protein-centric $F_{\max}$ , $S_{\min}$ , and AUPR, and the class-centric average AUC.

By embedding proteins with ESM2 [9] and employing graph attention mechanisms, our model further enriched the protein feature with contextual information present in the PPI network. However, the results indicated that incorporating PPIs as background information did not improve function prediction in our case. Upon scrutinizing the interaction data, we noticed that the interaction information was excessively sparse, failing to provide substantial support for function prediction and, instead, introducing additional noise. Our datasets included 10,107 proteins and 14,524 interactions, but only 1,935 proteins had interactions. Given the sub-optimal performance and sparse nature of PPI data, we excluded the DeepGOMeta-PPI model from further evaluation.

### Evaluation and comparison on the time-based split

We used a time-based split to evaluate DeepGOMeta as microbial data often contains an abundance of novel proteins. This is to ensure that our model is robust and effective in predicting the functions of these newly discovered proteins. We did this by comparing DeepGOMeta predictions on the newly annotated proteins with other state-of-the-art methods that predict functions based on protein language model embeddings and transformer-based deep learning models, including TALE [17], SPROF-GO [18] and DeepGO-SE [33]. We found that DeepGOMeta outperforms the DeepGO-SE method in all three sub-ontology evaluations and performs better than all the compared methods in the BPO and CCO evaluations in terms of *F*_max_ and *S*_min_. However, it resulted in lower performance than SPROF-GO method in the MFO evaluation and in terms of AUC in BPO evaluation. Table 6 shows the results of this evaluation.

**Table 6.** Evaluation of DeepGOMeta on time-based split.

| Method | MFO |  |  | BPO |  |  | CCO |  |  |
| --- | --- | --- | --- | --- | --- | --- | --- | --- | --- |
| | $F_{\max}$ | $S_{\min}$ | AUC | $F_{\max}$ | $S_{\min}$ | AUC | $F_{\max}$ | $S_{\min}$ | AUC |
| DeepGO-SE | 0.369 | 11.569 | 0.613 | 0.371 | 19.704 | 0.592 | 0.696 | 5.252 | 0.713 |
| SPROF-GO | **0.383** | **11.299** | **0.740** | 0.422 | 19.035 | **0.718** | 0.661 | 5.427 | 0.750 |
| TALE | 0.229 | 13.332 | 0.662 | 0.283 | 22.687 | 0.630 | 0.653 | 6.579 | 0.620 |
| DeepGOMeta | 0.371 | 11.967 | 0.602 | **0.431** | **18.296** | 0.644 | **0.749** | **4.497** | **0.758** |

### Evaluation strategies on microbial proteins

Given the unique challenges presented by microbial data and the lack of robust evaluation strategies, it was necessary to develop new strategies to assess the performance of DeepGOMeta in annotating microbial proteins in comparison with current annotation methods. Our evaluation employs k-means clustering and clustering purity based on true phenotype labels as a key metric (eq. 16) using a two-fold strategy. First, we compared our method against traditional sequence similarity-based methods by clustering based on sequence similarity. Second, we compared our method against database annotations by clustering based on semantic similarity.

Sequence similarity is a well-established method often employed in homology-based function prediction, and we aim to illustrate how DeepGOMeta performs in comparison to this approach. We used DeepGOMeta to annotate 2,000 proteins derived from microbial data in the MGnify database, and we calculated pairwise sequence similarity for these proteins. We clustered the proteins based on their sequence similarity scores and calculated purity, and in order to allow for an evaluation of our predicted functions against this, we clustered the proteins based on their predicted functions using semantic similarity. Both methods yielded a clustering purity of 0.55 (Figure 2). This implies that DeepGOMeta is at least as effective as traditional sequence similarity-based approaches, based on the assumption that a a similar degree of clustering purity based on the true phenotype labels indicates similar performance.

**Fig. 2.**
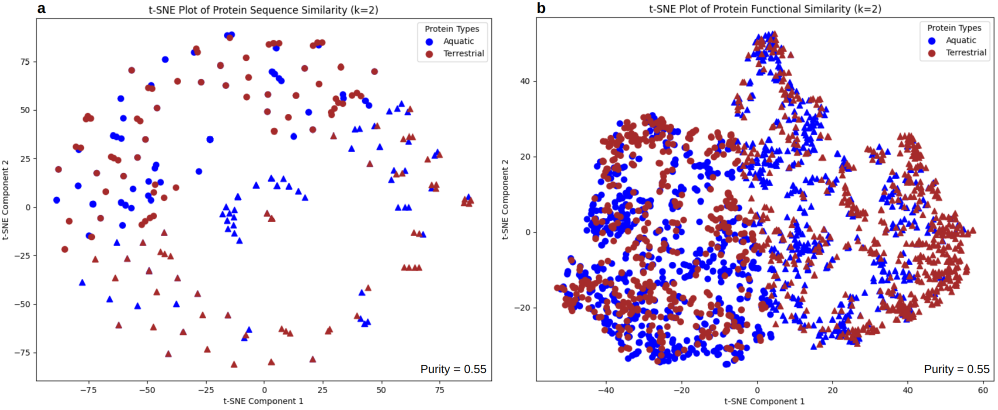
Clustering of microbial proteins (n = 2000) from MGnify. (a) Clustering based on sequence similarity between all proteins. (b) Clustering based on semantic similarity between all proteins.

As most current function annotation methods rely on annotations in existing databases, we subsetted this dataset to only keep proteins with Pfam annotations to compare against DeepGOMeta annotations. We observed that only 567 proteins have existing annotations, highlighting the annotation limitation in these traditional databases. DeepGOMeta was capable of annotating all 2,000 proteins, demonstrating its comprehensive annotation coverage. When focusing on the subset of 567 proteins with Pfam annotations, sequence similarity clustering yields a clustering purity of 0.6. After mapping the Pfam annotations to GO terms using Pfam2G), we found that the clustering purity using semantic similarity was also 0.6 for both Pfam and DeepGOMeta annotations (Figure 3). This parity in clustering purity might suggest that DeepGOMeta does not surpass sequence similarity methods in terms of predictive accuracy. However, it has the advantage that it can annotate all the proteins in the dataset.

**Fig. 3.**
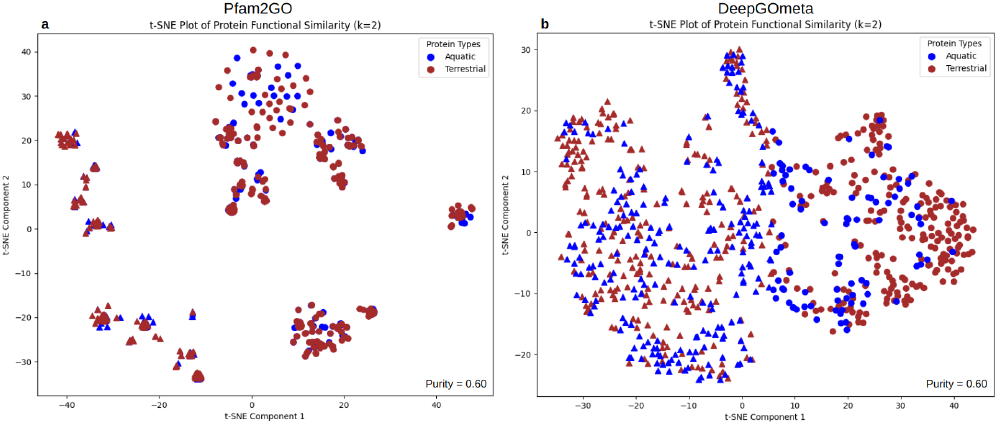
Clustering of microbial proteins (n = 567) from MGnify that possess Pfam annotations. (a) Clustering based on sequence similarity between all proteins. (b) Clustering based on semantic similarity between all proteins.

### Applications on amplicon and metagenome data

To demonstrate the utility of our method in function prediction for different types of microbial data, we used paired datasets of 16S amplicon reads and WGS reads of the same samples. Here, we employed our evaluation strategy where we used clustering to assess our method’s efficacy in capturing functionally relevant information from microbial communities. Due to the absence of ground-truth data for microbial functions, we assume that protein functions found in microbial communities are more similar when the microbial communities are from the same environment or share identical phenotypes. Consequently, we used functional similarity, based on the functions predicted by DeepGOMeta, to cluster microbial samples. This clustering, based on functional similarity, serves as an unsupervised and ostensibly unbiased method to group microbial communities by their functions. This approach allowed us to explore the primary drivers of community composition, focusing on the application of DeepGOMeta for gaining biological insights.

We used DeepGOMeta to construct functional profiles for each sample using reads from both sequencing strategies and compared against taxonomy-based clustering (Table 7). For each dataset, based on DeepGOMeta results, we constructed a binary representation of functions which indicates presence or absence of a function. For 16S data, we also constructed an abundance-weighted matrix, in which each function is assigned a weight (eq. 17). In certain contexts, DeepGOMeta demonstrated superior performance over OTU-based clustering. Specifically, in 5 out of the 9 phenotypes we analyzed, employing 16S functions (abundance-weighted) proved to be either on par with or more effective than clustering by OTUs. This suggests that DeepGOMeta’s functional profiles can be effective in capturing specific functional attributes that are unique to each phenotype. In some datasets, such as Mammalian Stool and Cameroon (Region, Ethnicity), the functional attributes were more defining than taxonomic composition, suggesting that these community compositions are driven by functions (in contrast to taxa).

**Table 7.** K-means cluster purity based on genus abundance and function abundance (k = number of true phenotype labels) - Transposed.

| Datatype | Mammalian Stool | Blueberry |  | India |  | Cameroon |  |  |  |
| --- | --- | --- | --- | --- | --- | --- | --- | --- | --- |
|  | Host | Type | Location | Location | Diet | Region | Diet | Sex | Ethnicity |
| OTUs | 0.63 | **0.63** | **0.77** | **0.62** | **0.73** | 0.41 | 0.43 | **0.61** | 0.67 |
| 16S Functions (Abundance) | **0.72** | **0.63** | 0.68 | 0.58 | **0.73** | **0.44** | 0.43 | 0.56 | **0.69** |
| 16S Functions (Binary) | 0.64 | 0.55 | 0.68 | 0.58 | **0.73** | **0.44** | 0.48 | 0.57 | 0.67 |
| WGS Functions | 0.55 | 0.55 | 0.68 | 0.58 | **0.73** | 0.41 | **0.52** | 0.56 | **0.69** |
| Pathways | 0.60 | 0.61 | 0.67 | **0.62** | 0.69 | 0.38 | 0.48 | 0.55 | 0.64 |

Conversely, in 3 out of 9 phenotypes studies, OTU-based clustering proved more effective. Specifically, in two datasets (Blueberry, India), the location phenotype was better explained by OTU composition than by functions. Interestingly, we found that using 16S functions in a binary format never outperformed the abundance-weighted approach, suggesting its limited efficacy. In the case of WGS functions, this method only took the lead in 1 out of 9 phenotypes, possibly indicating the necessity of weighing functions.

We also compared OTU-based clustering and DeepGOMeta-derived functional profiles with pathways generated by PICRUSt2 (detailed in the methods section). PICRUSt2 provides functional insights into microbial communities through KEGG/MetaCyc pathways. In only one case, the Cameroon dataset (Diet), we find that the functional insights provided by PICRUSt2 exhibit a better capacity to separate the phenotype than taxonomy-based clustering. We also find that pathway predictions and taxonomic composition separate samples by location equally well in the India dataset. However, for other datasets, there is no clear distinction between pathway-based and taxonomy-based clustering purity; none of which show a clear superiority in separating the samples between the phenotypes.

Compared to DeepGOMeta, PICRUSt2’s pathway information would be considered limited, as it constitutes only a subset of the predictable functions by DeepGOMeta in the form of BPO predictions. The results also indicate that PICRUSt2’s functional information overall does not separate samples better, based on phenotype, in comparison to DeepGOMeta. However, the experiment falls short of comparing the performance of the two function prediction methods. This indicates either a lack of strong associations between pathways and phenotypes or limitations of the algorithm/database used by PICRUSt2.

## Discussion

In this study we introduced DeepGOMeta, which aims to overcome the limitations of current methods in their lack of representative training sets and the lack of validation and applications on microbial data. Current function prediction methods are predominantly trained on eukaryotic data. We trained, tested, and evaluated three different models on UniProtKB/Swiss-Prot Knowledgebase proteins that belong to microbial species (prokaryotes, archaea, viral), a set more representative of species prevalent in microbial datasets. DeepGOMeta provides function predictions in the form of GO terms, as each of the three models was trained on a distinct GO sub-ontology. DeepGOMeta demonstrates an improvement over similarity-based benchmark methods in most evaluation metrics across the three sub-ontologies. In the comparison using a time-based split, DeepGOMeta outperformed DeepGO-SE, TALE and SPROF-GO in all three sub-ontology evaluations in BPO and CCO assessments in *F* max and *S*min metrics. However, in the MFO evaluation, the model was outperformed by SPROF-GO.

To evaluate the method’s predictions on microbial proteins, we designed a novel evaluation and benchmark strategy in which we use k-means clustering and clustering purity based on true phenotype labels in order to evaluate our method against sequence similarity-based methods and annotations in existing databases. For this, we use both sequence similarity-based clustering and semantic similarity-based clustering. We demonstrated that DeepGOMeta performs as well as traditional sequence similarity approaches in annotating 2,000 proteins from the MGnify protein database. This indicates the method’s ability to group proteins based on the environment in which they were found based on their predicted function. Notably, while only 567 proteins had existing Pfam annotations, DeepGOMeta successfully annotated all 2,000 proteins, showcasing its comprehensive annotation capabilities. While DeepGOMeta successfully annotated these proteins, extending the functional knowledge base, we recognize that further validation is necessary to ensure the specificity and accuracy of these annotations. The absence of Pfam annotations for this subset presents a challenge in directly validating the functional relevance and precision of our model’s predictions.

We demonstrated an application of DeepGOMeta in annotating both amplicon and metagenomic data in diverse datasets using a workflow that we have developed for this purpose. We constructed functional profiles for each samples based on 16S amplicon and WGS data, and compared the clustering of phenotypes against clustering based on taxonomic classification, allowing us to explore the primary drivers of community composition. We found that, overall, generating functional profiles using 16S amplicon data with DeepGOMeta (abundance-weighted) yields a higher clustering purity than clustering by OTUs. This indicates that many phenotypic differences can be attributed to functional variability rather than taxonomic composition. However, the variability in performance across the different datasets and phenotypes highlights that microbial community composition could be driven by functions and/or taxonomy. Looking forward, we propose several avenues for further enhancing the method’s utility. We plan to expand the training data to incorporate eukaryotic microbial genomes for WGS analysis, which will enable a more comprehensive understanding of metagenomic samples in ecosystems where eukaryotes play significant roles. Furthermore, as our observations reveal the sparsity of PPIs in bacteria, we intend to incorporate methods for interaction predictions. Integrating such methods as features within DeepGOMeta could substantially enhance its predictive accuracy and, consequently, our understanding of microbial interactions.

Additionally, we plan to expand our bioinformatics workflow in two ways. First, we aim to explore several ways through which we can assign weights to functions assigned to proteins from WGS data. Second, we would use the predicted functions and interactions to elucidate pathways both within and between organisms. This shift towards unraveling more complex biological processes will facilitate a deeper understanding of the intricate interactions and dependencies within microbial communities.

## Competing interests

No competing interest is declared.

## Acknowledgments

This work has been supported by funding from King Abdullah University of Science and Technology (KAUST) Office of Sponsored Research (OSR) under Award No. URF/1/4355-01-01, URF/1/4675-01-01, URF/1/4697-01-01, URF/1/5041-01-01, REI/1/5334-01-01, FCC/1/1976-46-01, and FCC/1/1976-34-01. This work was supported by the SDAIA-KAUST Center of Excellence in Data Science and Artificial Intelligence (SDAIA-KAUST AI). We acknowledge support from the KAUST Supercomputing Laboratory.

## Footnotes

1 https://github.com/bio-ontology-research-group/16SProcessing

## Notes

https://github.com/bio-ontology-research-group/deepgometa

### Competing Interest Statement

The authors have declared no competing interest.

### Summary of Updates

Corrected typo in the title and corrected the order of the authors

## References

1. Claudio Mirabello and B. Wallner. rawmsa: End-to-end deep learning using raw multiple sequence alignments. PLoS ONE, 14, 2019.

2. Jack A Gilbert, Dawn Field, Paul Swift, Simon Thomas, Denise Cummings, Ben Temperton, Karen Weynberg, Susan Huse, Margaret Hughes, Ian Joint, Paul J Somerfield, and Martin Mühling. The taxonomic and functional diversity of microbes at a temperate coastal site: a ‘multi-omic’ study of seasonal and diel temporal variation. PloS one, 5(11):e15545, 2010.

3. Maffei V.J. Zaneveld J.R. et al. Douglas, G.M. Picrust2 for prediction of metagenome functions. Nature Biotechnology, 38:685–688, 2020.

4. Franziska Wemheuer, Jessica A Taylor, Rolf Daniel, Eric Johnston, Peter Meinicke, Torsten Thomas, and Bernd Wemheuer. Tax4fun2: prediction of habitat-specific functional profiles and functional redundancy based on 16s rrna gene sequences. Environmental Microbiomes, 15(1):11, 2020.

5. S. F. Altschul, T. L. Madden, A. A. Schäffer, J. Zhang, Z. Zhang, W. Miller, and D. J. Lipman. Gapped BLAST and PSI-BLAST: a new generation of protein database search programs. Nucleic acids research, 25(17):3389–3402, September 1997.

6. Predrag Radivojac and et al. A large-scale evaluation of computational protein function prediction. Nat Meth, 10(3):221–227, January 2013. [PubMed:23353650] [PubMed Central:PMC3584181] [doi:10.1038/nmeth.2340].

7. M.V. Vecherskii, M.V. Semenov, A.A. Lisenkova, et al. Metagenomics: A new direction in ecology. Biology Bulletin Reviews, 48(Suppl 3):S107–S117, 2021.

8. M Mahmud, MS Kaiser, TM McGinnity, and A Hussain. Deep learning in mining biological data. Cognitive Computation, 13(1):1–33, 2021.

9. Zeming Lin, Halil Akin, Roshan Rao, Brian Hie, Zhongkai Zhu, Wenting Lu, Nikita Smetanin, Robert Verkuil, Ori Kabeli, Yaniv Shmueli, Allan dos Santos Costa, Maryam Fazel-Zarandi, Tom Sercu, Salvatore Candido, and Alexander Rives. Evolutionary-scale prediction of atomic-level protein structure with a language model. Science, 379(6637):1123–1130, 2023.

10. The UniProt Consortium. UniProt: the Universal Protein Knowledgebase in 2023. Nucleic Acids Research, 51(D1):D523–D531, 11 2022.

11. Benjamin Buchfink, Chao Xie, and Daniel H Huson. Diamond: a fast and sensitive alignment tool for shotgun metagenomic data. Genome research, 25(11):1755–1761, 2015.

12. The Gene Ontology Consortium. The gene ontology resource: 20 years and still going strong. Nucleic Acids Research, 47(D1):D330–D338, 2019.

13. Damian Szklarczyk and et al. The string database in 2019: quality-controlled protein–protein association networks, made broadly accessible. Nucleic acids research, 47(D1):D607–D613, 2019.

14. LJ Richardson and et al. Mgnify: the microbiome sequence data analysis resource in 2023. Nucleic Acids Research, 2023.

15. F Meyer, D Paarmann, M D’Souza, R Olson, EM Glass, M Kubal, T Paczian, A Rodriguez, R Stevens, A Wilke, J Wilkening, and RA Edwards. The metagenomics rast server – a public resource for the automatic phylogenetic and functional analysis of metagenomes. BMC Bioinformatics, 9:386, 2008.

16. ER Morton, J Lynch, A Froment, S Lafosse, E Heyer, M Przeworski, R Blekhman, and L Ségurel. Variation in rural african gut microbiota is strongly correlated with colonization by entamoeba and subsistence. PLoS Genet, 11(11):e1005658, Nov 2015.

17. Yue Cao and Yang Shen. TALE: Transformer-based protein function Annotation with joint sequence–Label Embedding. Bioinformatics, 37(18):2825–2833, 03 2021.

18. Qianmu Yuan, Junjie Xie, Jiancong Xie, Huiying Zhao, and Yuedong Yang. Fast and accurate protein function prediction from sequence through pretrained language model and homology-based label diffusion. Briefings in Bioinformatics, 24(3):bbad117, 03 2023.

19. Shaojun Wang, Ronghui You, Yunjia Liu, Yi Xiong, and Shanfeng Zhu. Netgo 3.0: Protein language model improves large-scale functional annotations. Genomics, Proteomics & Bioinformatics, 21(2):349–358, 2023.

20. Maxat Kulmanov and Robert Hoehndorf. DeepGOPlus: improved protein function prediction from sequence. Bioinformatics, 07 2019. [PubMed:31350877] [doi:10.1093/bioinformatics/btz595].

21. Maxat Kulmanov and Robert Hoehndorf. DeepGOZero: improving protein function prediction from sequence and zero-shot learning based on ontology axioms. Bioinformatics, 38(Supplement 1):i238–i245, 06 2022.

22. Maxat Kulmanov, Wang Liu-Wei, Yuan Yan, and Robert Hoehndorf. El embeddings: Geometric construction of models for the description logic el++. In Proceedings of the Twenty-Eighth International Joint Conference on Artificial Intelligence, IJCAI-19, pages 6103–6109. International Joint Conferences on Artificial Intelligence Organization, 7 2019.

23. Ahmed Elnaggar and et al. Prottrans: Toward understanding the language of life through self-supervised learning. IEEE Transactions on Pattern Analysis and Machine Intelligence, 44(10):7112–7127, 2022.

24. Ron Caspi, Richard Billington, Ingrid M Keseler, Anamika Kothari, Markus Krummenacker, Peter E Midford, Wai Kit Ong, Suzanne Paley, Pallavi Subhraveti, and Peter D Karp. The MetaCyc database of metabolic pathways and enzymes - a 2019 update. Nucleic Acids Research, 48(D1):D445–D453, 10 2019.

25. Philip Resnik. Semantic similarity in a taxonomy: An information-based measure and its application to problems of ambiguity in natural language. Journal of artificial intelligence research, 11:95–130, 1999.

26. Robert D Finn, Penny Coggill, Ruth Y Eberhardt, Sean R Eddy, Jaina Mistry, Alex L Mitchell, Simon C Potter, Marco Punta, Matloob Qureshi, Amaia Sangrador-Vegas, et al. The pfam protein families database. Nucleic acids research, 42(D1):D222–D230, 2014.

27. K Clark, I Karsch-Mizrachi, DJ Lipman, J Ostell, and EW Sayers. Genbank. Nucleic Acids Research, 44(D1):D67–D72, 2016.

28. N. O’Leary, MW Wright, JR Brister, S Ciufo, D Haddad, R McVeigh, B Rajput, B Robbertse, B Smith-White, D Ako-Adjei, et al. Reference sequence (refseq) database at ncbi: current status, taxonomic expansion, and functional annotation. Nucleic Acids Research, 44(D1):D733–D745, 2016.

29. Shifu Chen, Yanqing Zhou, Yaru Chen, and Jia Gu. fastp: an ultra-fast all-in-one fastq preprocessor. Bioinformatics, 34(17):i884–i890, 2018.

30. Ben Langmead and Steven L Salzberg. Fast gapped-read alignment with bowtie 2. Nature Methods, 9(4):357–359, 2012.

31. Dinghua Li, Chi-Ming Liu, Ruibang Luo, Kunihiko Sadakane, and Tak-Wah Lam. Megahit: an ultra-fast single-node solution for large and complex metagenomics assembly via succinct de bruijn graph. Bioinformatics, 31(10):1674–1676, 2015.

32. Doug Hyatt, Gwo-Liang Chen, Philip F Locascio, Miriam L Land, Frank W Larimer, and Loren J Hauser. Prodigal: prokaryotic gene recognition and translation initiation site identification. BMC bioinformatics, 11(1):119, 2010.

33. Maxat Kulmanov, Francisco J. Guzmán-Vega, Paula Duek Roggli, Lydie Lane, Stefan T. Arold, and Robert Hoehndorf. Deepgo-se: Protein function prediction as approximate semantic entailment. bioRxiv, 2023.

